# Containing Emerging Epidemics: a Quantitative Comparison of Quarantine and Symptom Monitoring

**DOI:** 10.1101/072652

**Authors:** Corey M Peak, Lauren M Childs, Yonatan H Grad, Caroline O Buckee

## Abstract

Strategies for containing an emerging infectious disease outbreak must be non-pharmaceutical when drugs or vaccines for the pathogen do not yet exist or are unavailable. The success of these non-pharmaceutical strategies will depend not only on the effectiveness of quarantine or other isolation measures but also on the epidemiological characteristics of the infection. However, there is currently no systematic framework to assess the relationship between different containment strategies and the natural history and epidemiological dynamics of the pathogen. Here, we compare the effectiveness of quarantine and symptom monitoring, implemented via contact tracing, in controlling epidemics using an agent-based branching model. We examine the relationship between epidemic containment and the disease dynamics of symptoms and infectiousness for seven case study diseases with diverse natural histories including Ebola, Influenza A, and Severe Acute Respiratory Syndrome (SARS). We show that the comparative effectiveness of symptom monitoring and quarantine depends critically on the natural history of the infectious disease, its inherent transmissibility, and the intervention feasibility in the particular healthcare setting. The benefit of quarantine over symptom monitoring is generally maximized for fast-course diseases, but we show the conditions under which symptom monitoring alone can control certain outbreaks. This quantitative framework can guide policy-makers on how best to use non-pharmaceutical interventions to contain emerging outbreaks and prioritize research during an outbreak of a novel pathogen.

**SIGNIFICANCE:** Quarantine and symptom monitoring of contacts with suspected exposure to an infectious disease are key interventions for the control of emerging epidemics; however, there does not yet exist a quantitative framework for comparing the control performance of each. Here, we use a mathematical model of seven case study diseases to show how the choice of intervention is influenced by the natural history of the infectious disease, its inherent transmissibility, and the intervention feasibility in the particular healthcare setting. We use this information to identify the most important characteristics of the disease and setting that need to be characterized for an emerging pathogen in order to make an informed decision between quarantine and symptom monitoring.

## INTRODUCTION

The global burden of emerging infectious diseases is growing and prompts the need for effective containment policies (1–3). In many cases, strategies must be non-pharmaceutical, as targeted drugs or vaccines for the pathogens are unavailable. Among the various containment strategies, isolation of ill and potentially infectious patients is one of the most intuitive, relying on the tracing the contacts of known cases. Contacts with symptoms can then be hospitalized or isolated, but policy makers must also decide how best to handle contacts that do not meet the case definition for infection. Two strategies have historically been used in the case of a potentially infected but symptom-free contact: quarantine and symptom monitoring.

Quarantine of potentially infected contacts during an epidemic is highly conservative with respect to efficacy, but it comes at a high cost. Costs associated with quarantine policies range from direct (e.g., implementation expenses and the restriction of personal liberties) to indirect (e.g., stigmatization of health workers and sometimes interruption of financial and trade markets) (4–8). A less conservative but substantially cheaper and more socially palatable approach is active symptom monitoring of contacts. In this strategy, health workers check on contacts one or two times a day and isolate them if symptoms occur (see definitions in *Methods*).

Given the importance of rapid decision making in the event of novel emerging pathogens, and the potentially devastating consequences of poor containment strategies, quantitative guidelines are urgently needed for deciding whether quarantine is, according to Gates et al., at worst “counterproductive” or at best “one of the few tactics that can reduce its spread” (9). Current guidance on the use of quarantine or symptom monitoring is *ad hoc* and disease-specific, lacking the generalizability required for rapid decision-making for novel pathogens and leading to confusion during implementation (10–12). During the Severe Acute Respiratory (SARS) epidemic, broad quarantine interventions were applied in Taiwan and subsequently abandoned (13). Furthermore, we are aware of no framework that considers the implementation setting as a factor for intervention choice or performance, despite its obvious importance. Indeed, the United States Centers for Disease Control and Prevention (CDC) implicitly recognized the value of implementation setting by differentiating its international response, where quarantine was prioritized (14), and its domestic response, where symptom monitoring was prioritized (15,16).

The success of these approaches is not simply a reflection of the efficiency of their implementation but crucially depend on the biology and natural history of the pathogen in question. Previous theoretical work by Fraser, *et al*. (17) summarized these dynamics into a measure of the proportion of infections by asymptomatic infection (*θ*) and the basic reproductive number (*R*_0_), defined as the average number of infections generated by an infectious individual in a fully susceptible population. Subsequent work has explored the interaction between disease characteristics (e.g., super-spreading (18)) and the performance of interventions (e.g., travel screening (19)), but the recent Ebola epidemic demonstrated that at least two large questions remain (7). Firstly, what is the role of symptom monitoring as an alternative to quarantine, and secondly, how does this choice depend on the characteristics of the disease, the setting, and their interactions?

Here we develop an agent-based branching model that accommodates realistic distributions of disease characteristics and maintain the infector-infectee correlation structure necessary for interventions targeted via contact tracing. To assess diseases with a wide range of natural histories that have the potential for causing sudden, severe epidemics, we consider case studies of seven known pathogens: Ebola; hepatitis A; influenza A; Middle East Respiratory Syndrome (MERS); pertussis; SARS; and smallpox. We identify which disease characteristics and intervention attributes are most critical in deciding between quarantine and symptom monitoring, and provide a clear, general framework for understanding the consequences of isolation policies during an epidemic of known or novel pathogens.

## RESULTS

### Intervention Effectiveness Depends on Disease Epidemiological Dynamics

To assess the impact of quarantine and symptom monitoring, we developed a general mathematical model of disease transmission and interventions targeted via contact tracing (**Fig 1**). The model structure accommodates five key metrics of intervention performance in a given setting (**Table 1**). We used particle filtering to generate parameter sets consistent with seven case studies of outbreak-prone pathogens (see *Methods* and (**Table 2**).

**Fig. 1.**
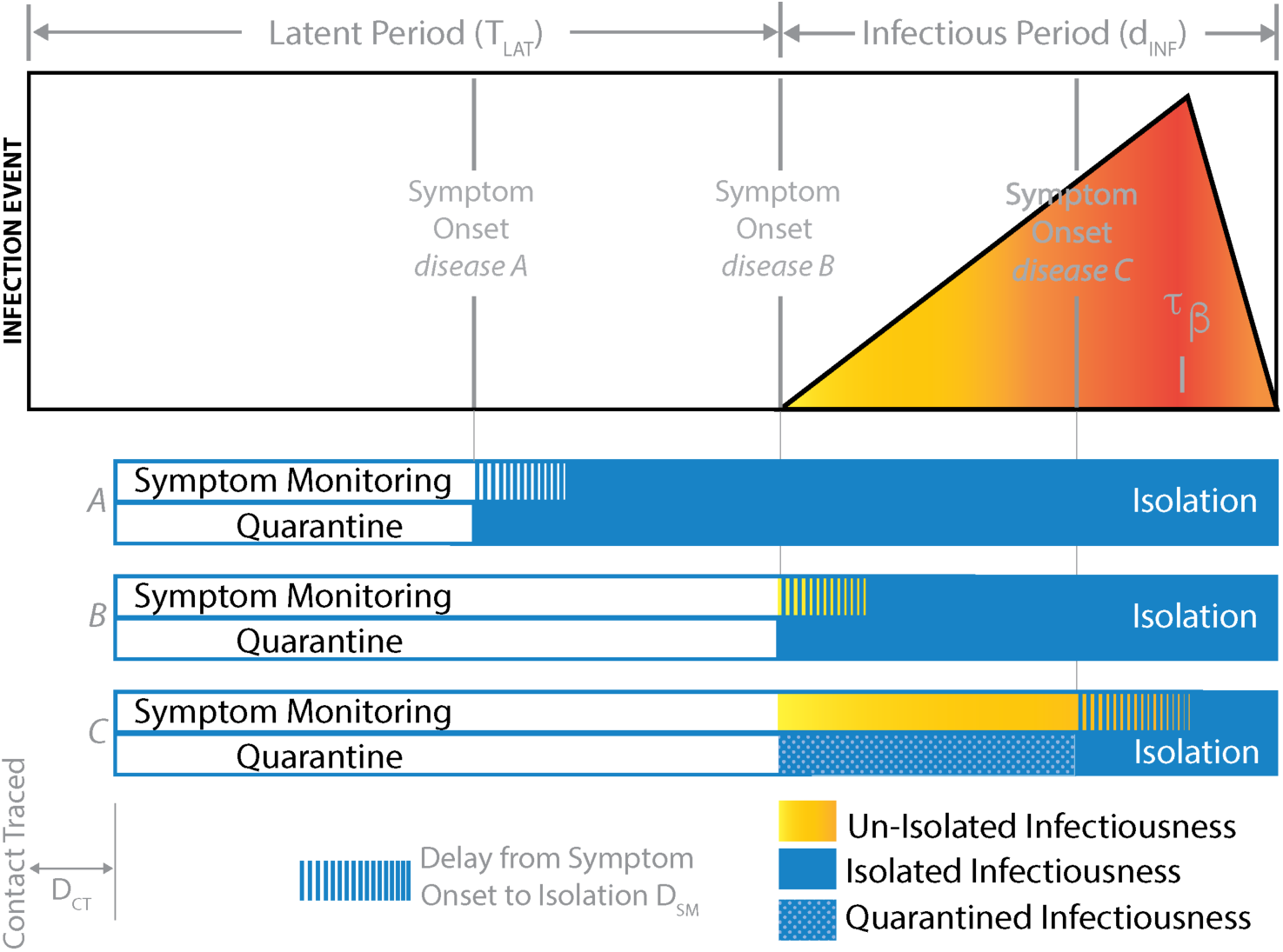
Schematic of the natural history of disease and the timing of interventions. Beginning on the left with the infection event, one progress through a latent period (T_LAT_) before becoming infectious for d_INF_ days with late peak infectiousness τ_*β*_. For diseases A, B, and C, symptoms are respectively shown to emerge before, concurrent with, and after onset of infectiousness. We show here an individual who is traced shortly after infection and is placed under symptom monitoring or quarantine after a short delay D_CT_.

**Table 1.** Intervention Parameters.

|  | Variable | Example Intervention Performance |  |
| --- | --- | --- | --- |
|  |  | Optimal | High |
| Isolation Effectiveness | $\gamma$ | 1 | 0.9 |
| Fraction of contacts traced | $P_{CT}$ | 1 | 0.9 |
| Fraction of traced contacts who are truly infected | $P_{INF}$ | 1 | 0.5 |
| Delay in tracing a named contact | $D_{CT}$ | 0.25 days<br>$\pm 0.25$ | 0.5 days<br>$\pm 0.5$ |
| Delay from symptom onset to isolation | $D_{SM}$ | 0.25 days<br>$\pm 0.25$ | 0.5 days<br>$\pm 0.5$ |
| Delay from symptom onset to health seeking behavior | $D_{HSB}$ | Disease-dependent.<br>$\sim \text{unif}(0, d_{INF})$ * | |

\* The mean $D_{HSB}$ will equal the mean $d_{INF}$ reported in **Table 2**.

**Table 2.** Disease parameters

|  | Inputs from Published Estimates |  |  | Parameters fit via Sequential Monte-Carlo method |  |  |
| --- | --- | --- | --- | --- | --- | --- |
| | Basic Reproductive Number $R_0$ | Serial Interval (days) | Incubation Period $T_{INC}$ (days) | Latent period offset $T_{OFFSET} = T_{LAT} - T_{INC}$ (days) | Mean duration of infectiousness $(1 + d_{INF})/2$ (days) | Time of peak Infectiousness $\tau_\beta$ (range 0-1) |
| Ebola | 1.83 (34)<br>[1.72, 1.94] | 13.36 (35)<br>[2.66, 38.8] | 7.87 (36)<br>[0.93, 28.2] | 0.33<br>[0**, 1.01] | 6.53<br>[1.28, 13.7] | 0.10<br>[0, 0.37] |
| Hepatitis A | 2.25 *<br>[2, 2.5] | 26.72 (37)<br>[20.7, 33.8] | 29.11 (38)<br>[24.6, 34.1] | -5.33<br>[-7.57, -3.26] | 6.23<br>[1.22, 15.8] | 0.35<br>[0, 0.98] |
| Influenza A | 1.54 (39)<br>[1.28, 1.80] | 2.20 (37)<br>[0.63, 3.76] | 1.40 (40)<br>[0.63, 3.10] | -0.23<br>[-0.76, 0.29] | 1.88<br>[1.04, 3.84] | 0.49<br>[0.02, 0.98] |
| MERS | 0.95 (41)<br>[0.6, 1.3] | 7.62 (42)<br>[2.48, 23.3] | 5.20 (42)<br>[1.83, 14.7] | -1.55<br>[-3.14, 0.02] | 8.35<br>[1.37, 19.9] | 0.37<br>[0.01, 0.96] |
| Pertussis | 4.75 *<br>[4.5, 5] | 19.26 (43)<br>[3.61, 57.2] | 7.00 (33)<br>[4.00, 10.0] | -2.14<br>[-5.39, 0.78] | 34.38<br>[2.67, 76.7] | 0.45<br>[0.11, 0.88] |
| SARS | 2.9 (44)<br>[2.2, 3.6] | 8.32 (44)<br>[1.59, 19.2] | 4.01 (40)<br>[1.25, 12.8] | 0.16<br>[0**, 0.67] | 10.94<br>[1.50, 23.0] | 0.10<br>[0, 0.46] |
| Smallpox | 4.75 *<br>[4.5, 5] | 15.54 (45)<br>[9.98, 24.2] | 11.83 (45)<br>[8.47, 16.5] | 0.03<br>[-1.80, 1.68] | 8.45<br>[1.37, 20.0] | 0.32<br>[0, 0.97] |

Median (Reference); [95% CI]
\* Assumed. \*\* Sequential Monte-Carlo boundary condition reached

**Fig 2** illustrates model dynamics for two infections with different epidemiological characteristics, Ebola and influenza A, showing the range of epidemic outcomes resulting from different interventions for a given R_0_ value. Unimpeded exponential epidemic growth in our branching model (red) can be reduced by the increasingly conservative interventions of health-seeking behavior (teal), symptom monitoring (gold), and quarantine (blue) (**Figs 2A-B**). Under a given intervention policy, we estimate the effective reproductive number (R_e_) as the average number of infections generated by an infectious individual in the population. The effective reproductive number under symptom monitoring (R_S_), quarantine (R_q_), and the absolute difference between the two (*R_s_ – R_Q_*) increases with R_0_ and differs by disease (**Figs 2C-D**).

**Fig. 2.**
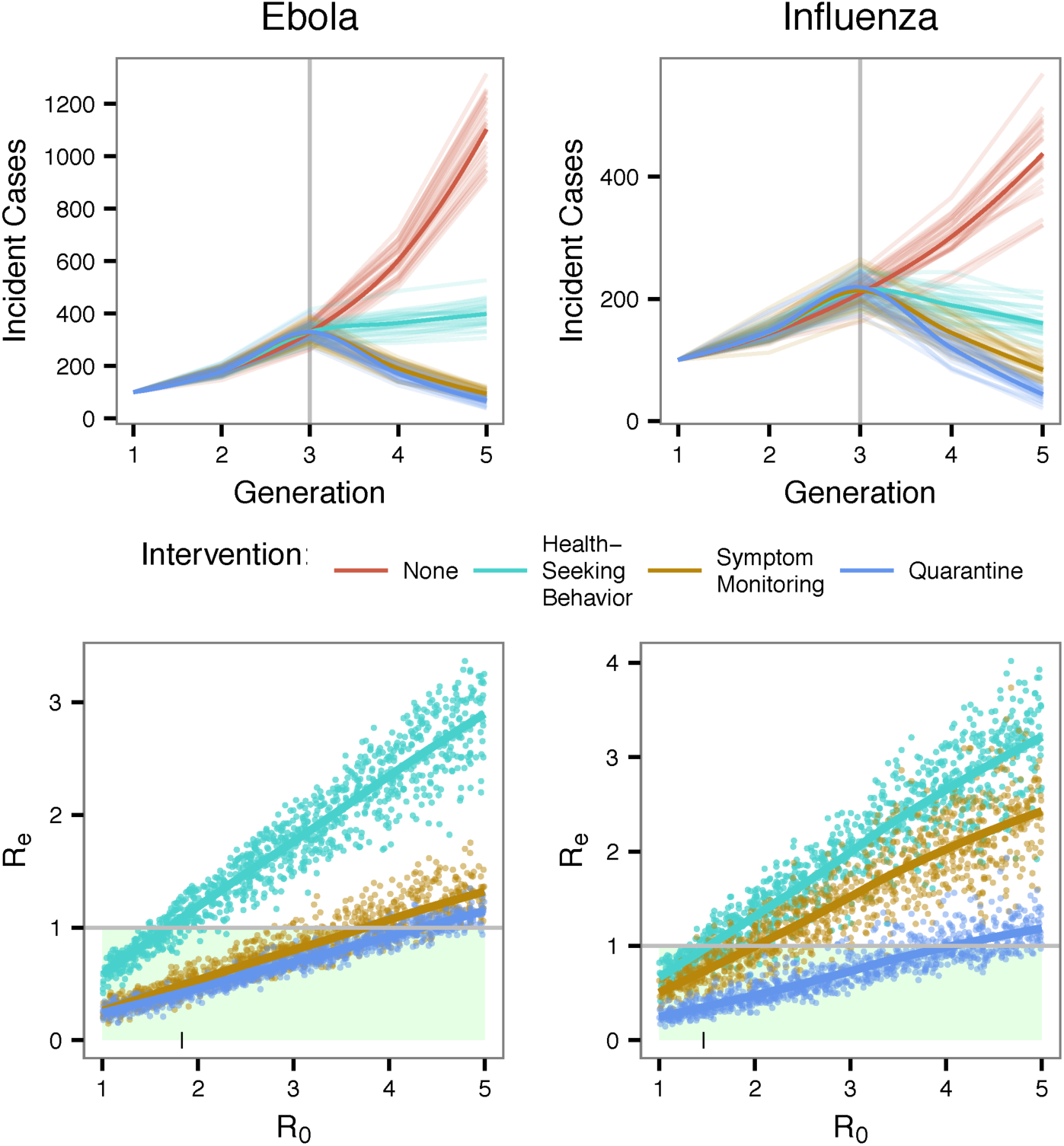
Model dynamics and output for two exemplar diseases: Ebola and influenza A. Each line in the panels (A) and (B) designates one model run initiated with 100 infectious individuals in Generation 1 and submitted to either no intervention (red), health seeking behavior (teal), symptom monitoring every day (gold), or quarantine (blue) at Generation 3. Each point in panels C and D designates the simulated effective reproductive number from one model run with input reproductive number (x-axis) between 1 and 5, with the small vertical line denoting the input R for panels (A) and (B) (1.83 and 1.46, respectively). Loess curves are shown as heavier lines. Here, symptom monitoring performs similarly to quarantine for Ebola control, but not for influenza A. Note the independent y-axes.

We find the effectiveness of symptom monitoring and quarantine in controlling a disease in a particular setting depends critically on its biological dynamics (e.g. latent and infectious periods) and transmissibility (*R*_0_) (**Fig 3a**). Holding transmissibility constant (*R*_0_ arbitrarily set to 2.75±0.25), biological dynamics alone strongly influence the effectiveness of quarantine and especially symptom monitoring, as seen by the wide spread in *R_s_* (**Fig 3b**).

**Fig. 3.**
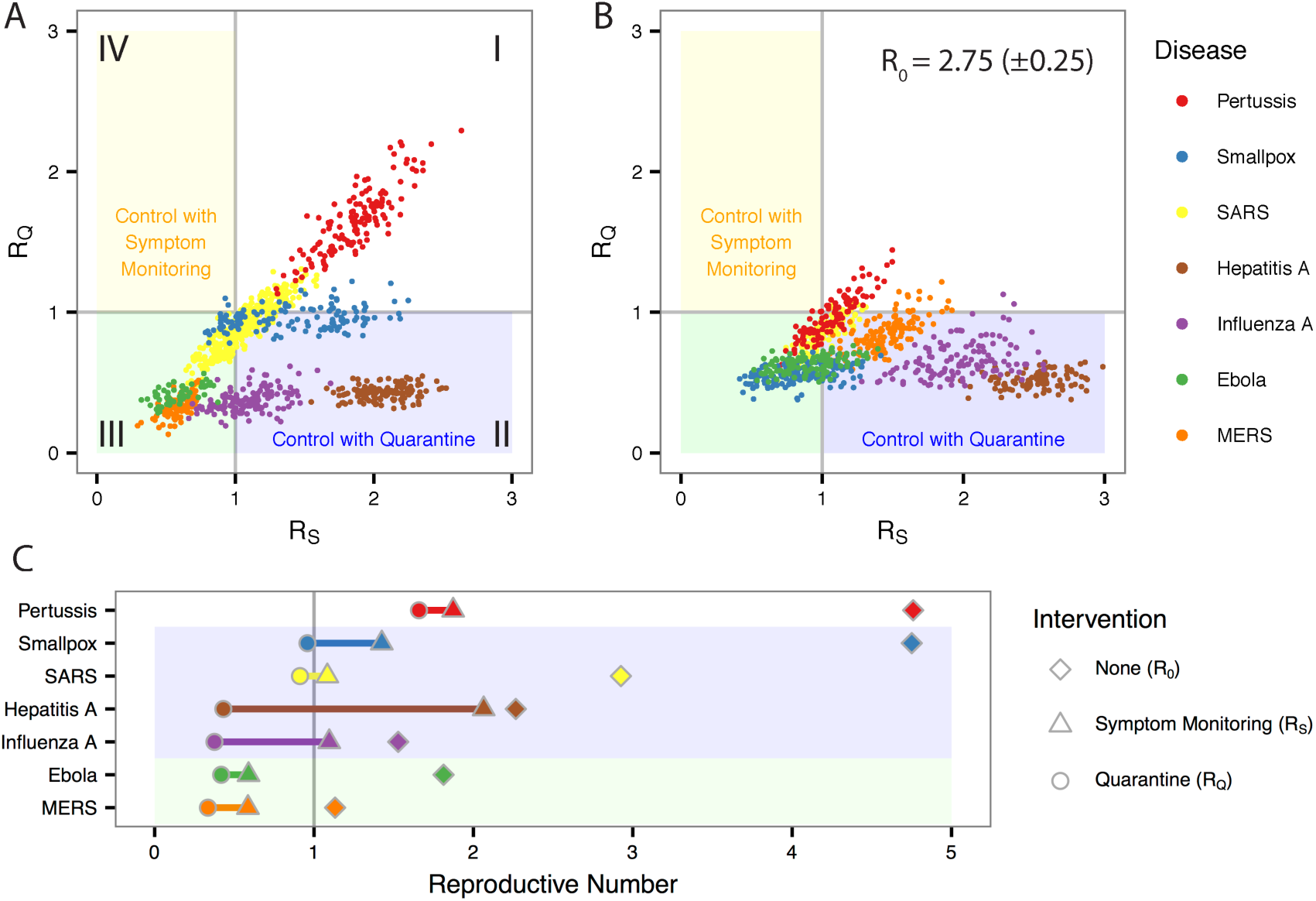
Infection control performance depends on disease biological dynamics and inherent transmissibility (R_0_). (A) The effective reproductive number under symptom monitoring (x-axis) and quarantine (y-axis) for 100 simulations of each disease when the basic reproductive number is set to published values [See Panel C, diamonds]. Quadrants indicate regions of control with (I) neither quarantine nor symptom monitoring, (II) only symptom monitoring, (III) either quarantine or symptom monitoring, and (IV) only quarantine. (B) As in (A), but the basic reproductive number (*R*_0_) is set to for all diseases to 2.75 (± 0.25) to isolate inherent differences in biological dynamics. (C) Disease-specific mean basic reproductive number (diamond) and the mean effective reproductive numbers under symptom monitoring (triangle) and quarantine (circle). The length of the horizontal line therefore equals the absolute comparative effectiveness *R_S_* – *R_Q_.*

In simulations with high intervention performance settings (i.e., 90% isolation effectiveness, 90% of contacts traced within one day, and symptoms monitored daily), diseases such as MERS and Ebola could be controlled (i.e., R_e_ < 1) with either quarantine or symptom monitoring; diseases such as hepatitis A with only quarantine; but diseases such as pertussis require additional interventions in order to reduce the effective reproductive number below one, due in large part to pre-symptomatic infectiousness (**Fig 3c**). The absolute comparative effectiveness (*R_s_ – R_Q_*) varies widely by disease, as demonstrated by the line length in **Fig 3c**. The relative comparative effectiveness 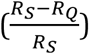 also varies widely, with quarantine reducing R_S_ by over 65% for influenza A and hepatitis A and by less than 10% for pertussis (**Fig S1**). The reader can explore results from landscapes with different intervention performance settings and disease transmissibility in the **interactive supplement** (https://coreypeak.shinyapps.io/InteractiveQuarantine).

### Categorizing Disease Control Frontiers

In order to compare the effectiveness of symptom monitoring and quarantine, one must select an appropriate metric to compare R_S_ and R_Q_. We therefore categorized intervention response heterogeneity into four control quadrants (**Fig 3a**). In quadrant I, where neither intervention is sufficient to prevent epidemic growth, the relative difference 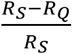 can distinguish whether quarantine is merited or could be paired with other strategies to achieve control. Because quarantine is by definition the more conservative intervention, simulation results in quadrant II occur only stochastically. In quadrant III, where both interventions are sufficient and the number of prevented cases can be more directly estimated, the distinguishing metric was the absolute difference *R_s_ – R_Q_* and its inverse 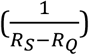, which can be interpreted as the number of contacts that must be quarantined in order to prevent one additional case over symptom monitoring (an analog of “number needed to treat”). For the example of SARS, Day et al. propose that mass quarantine may be unnecessary because effective symptomatic isolation alone would sufficiently control the disease (hence placing the disease in quadrant III) (8). In quadrant IV, where quarantine but not symptom monitoring can control the disease, quarantine would be strongly considered as the minimum sufficient strategy to prevent exponential epidemic growth.

The following two sections aim to identify which disease characteristics and intervention performance metrics most strongly influence these differences in response to quarantine and symptom monitoring.

### Ranking of epidemiological characteristics by importance for containment feasibility

The comparative effectiveness of quarantine and symptom monitoring is strongly influenced by differences in the infection's natural history. We measured partial rank correlation coefficients to examine which biological characteristics in particular are most influential after controlling for the other characteristics (*Methods*). As demonstrated by strongly negative partial rank correlation coefficients in **Fig 4**, increasing the duration of infectiousness (*d_INF_*) and elongating the latent period (*T_0FFSET_*) reduced the differences between quarantine and symptom monitoring, thereby making the interventions more similar. Other factors, such as overdispersed heterogeneity of the basic reproductive number (*κ*), did not influence the average effect of symptom monitoring and quarantine, as reflected by a coefficient of nearly zero. However, at a given effective reproductive number, overdispersion does decrease the average number of generations until extinction, as predicted (**Fig S2**) (18). Longer incubation periods (T_INC_) increased the preference for quarantine, as seen by the positive partial rank correlation coefficient for both absolute and relative comparative effectiveness. However, the length of the incubation period does not generally influence comparative effectiveness per quarantine day because the number of days in quarantine (*d_Q_*) increases as the incubation period lengthens (**Fig S3**).

**Fig. 4.**
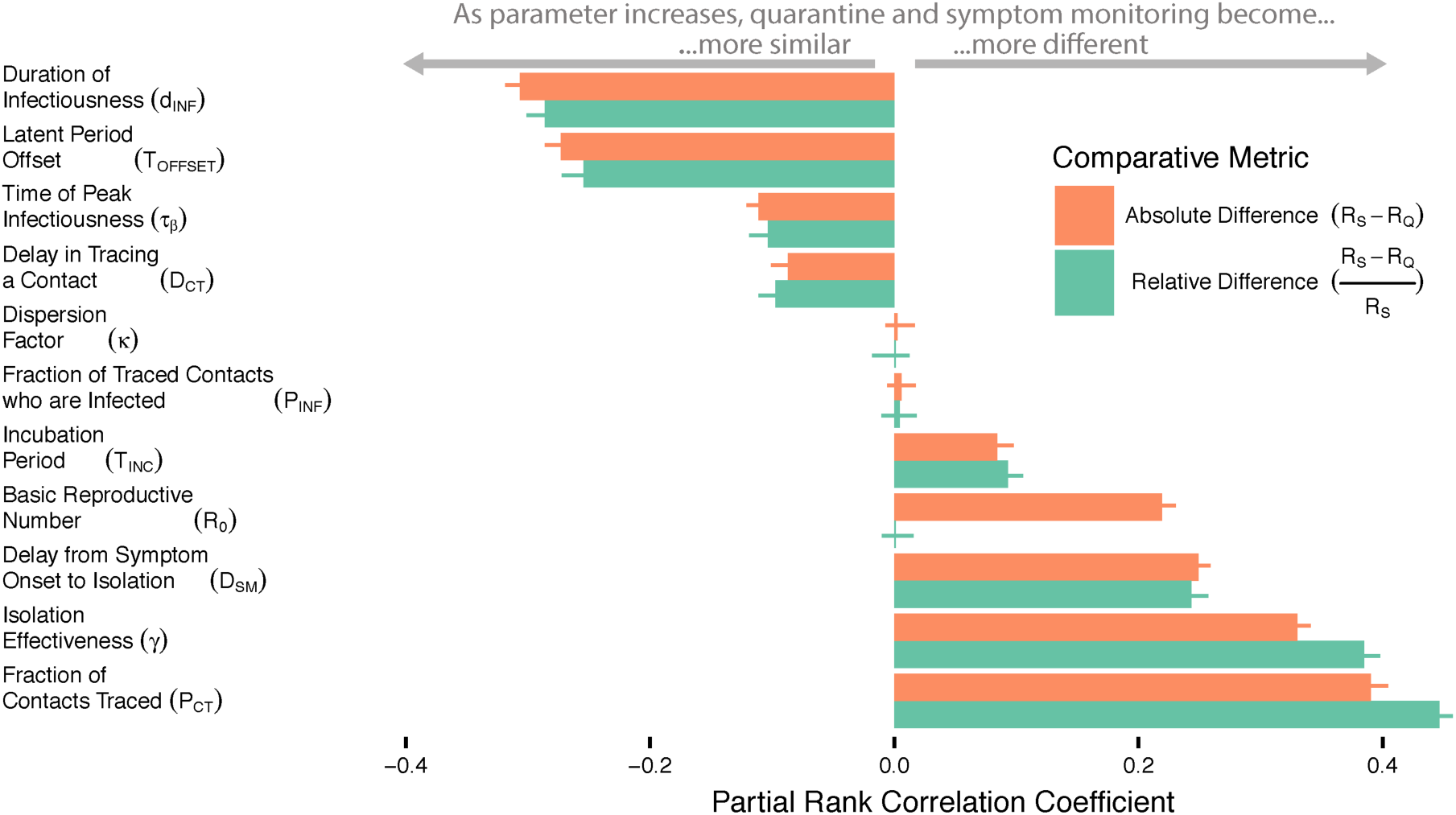
Influence of disease characteristics and intervention performance metrics. Partial rank correlation coefficients (x-axis) measuring the influence of disease characteristics and intervention performance metric (rows) on the absolute (red) and relative (green) comparative effectiveness of quarantine and symptom monitoring, pooled for all case study diseases. The 95% confidence intervals from 100 bootstrapped samples are represented by error bars.

Frequently, the most politically and economically pressing concerns are whether control (i.e. *R_e_* < 1) is logistically achievable and what would be the least invasive intervention to achieve control. **Fig 5** shows frontiers where control of an Ebola-like disease requires increasingly invasive interventions, moving from health-seeking behavior (teal), to symptom monitoring (gold), to quarantine (blue), the most invasive. **Fig 5a** shows how this frontier is influenced by the inherent transmissibility (*R*_0_) and timing of the latent period relative to the incubation period (*T_0FFSET_*), with all other characteristics similar to Ebola. When R_0_ is large and symptoms emerge long after infectiousness (e.g., *T_0FFSET_ >* 0), even quarantine is insufficient to control the disease with optimal intervention performance. However, we observe that when transmissibility is relatively low (e.g., *R*_0_ < 2.5), control of this hypothetical disease can be achieved even if infectiousness precedes symptoms by several days (**Fig 5a**) or if a substantial fraction of transmission events occur before symptom onset (adapting the framework of (17)) (**Fig 5b**).

**Fig. 5.**
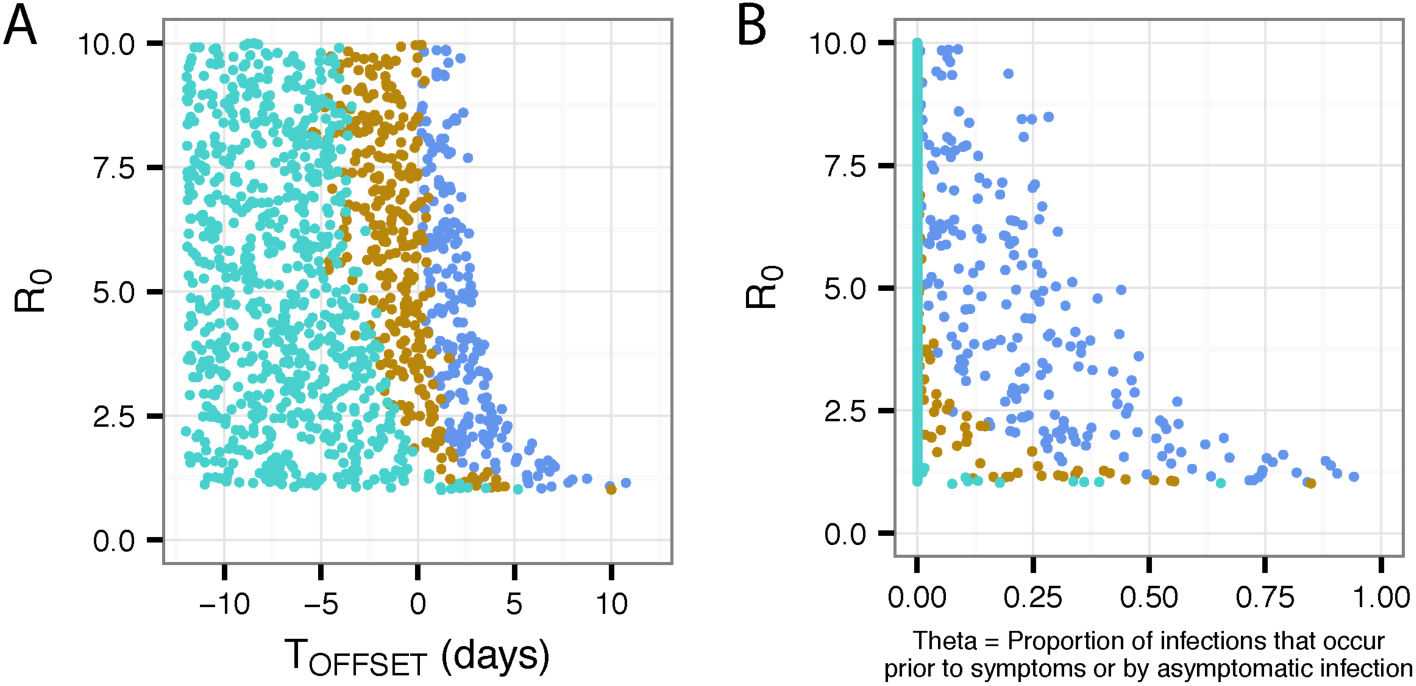
Minimally invasive interventions sufficient to control a hypothetical disease. (A) Disease characteristics drawn from Ebola except symptoms are assumed to either: precede infectiousness by up to 10 days (X = −10 days); coincide with infectiousness onset (X = 0 days); or emerge up to 10 days after infectiousness onset (X = +10 days). Points represent simulations where health-seeking behavior (teal), symptom monitoring (gold), or quarantine (blue) were the minimally sufficient intervention to bring *R_e_* below 1. (B) As in (A), but the x-axis is transformed to represent the proportion of infections that occur prior to symptoms in a analogous way to Fraser, Riley, et al 2004 (17) Interventions are in an optimal setting with all contacts being traced immediately, no infections occur during isolation, and symptom monitoring is performed twice per day.

### Ranking of intervention performance metrics by importance for containment feasibility

Policy makers facing an epidemic must also consider the expected performance of interventions, since the effectiveness of targeted control policies will depend on their feasibility within a particular healthcare system. Generally, we found the benefit of quarantine over symptom monitoring increases with better intervention performance (i.e. larger fraction of contacts traced (*P_CT_*), better isolation effectiveness (*γ*), and shorter delays in tracing a contact (*D_CT_*) (**Fig 4**). However, the effectiveness of symptom monitoring approached that of quarantine when the delay between symptom onset and isolation (*D_SM_*) is shortened, due either to more frequent symptom monitoring or more sensitive detection of symptoms followed by prompt isolation.

While these patterns were highly consistent across the case study diseases, some intervention performance metrics were particularly influential in the presence of certain disease characteristics. For example, diseases with short incubation periods (*T_1NC_*) such as influenza A were strongly influenced by delays in tracing a contact (*D_CT_*) (**Fig S3**).

## DISCUSSION

A key strategy to controlling the spread of infectious diseases focuses on tracing the contacts of infected individuals, with the goal of limiting subsequent spread should those contacts become infectious. Here, we present the first study comparing the effectiveness of the two primary non-pharmaceutical interventions targeted via contact tracing: symptom monitoring and quarantine. We show that the interventions are not equivalent and that the choice of which intervention to implement to achieve optimal control depends on the natural history of the infectious disease, its inherent transmissibility, and the intervention feasibility in the particular healthcare setting.

Our results show that the benefit of quarantine over symptom monitoring is maximized for fast-course diseases (short duration of infectiousness and a short latent period compared to the incubation period), and in settings where isolation is highly effective, a large fraction of contacts are traced, or when there is a long delay between symptom onset and isolation. This delay (*D_SM_*) not only captures ineffective symptom monitoring, but also the potential for symptoms to be masked for a period of time through biological (e.g., natural disease progression or self-medication with anti-pyretics) or behavioral (e.g., avoidance) mechanisms. In contrast, the widely-discussed “super spreading” disease characteristic did not independently impact the mean comparative effectiveness of interventions after holding R_0_ constant. However, this characteristic could remain important to understand disease control during highly stochastic stages of emergence and extinction (**Fig S2**) (18). Our findings are consistent with Fraser, Riley, et al. (17) that both inherent transmissibility and the proportion of transmission from asymptomatically infected individuals are key epidemiological parameters for the feasibility of control via quarantine.

In addition to identifying parameters that differentiate quarantine and symptom monitoring, our results also characterize parameter spaces where symptom monitoring, not just quarantine, is sufficient for containment of an emerging epidemic. Given the high costs and poor scalability of quarantine, symptom monitoring is likely to be a key intervention for future epidemic containment.

The CDC has current legal authority to quarantine for diseases including Ebola, SARS, MERS, smallpox, and influenza strains with pandemic potential (12). Our results support the retention of quarantine as a live-option for each of these diseases, but only if control is infeasible through symptom monitoring (i.e. *R_Q_* > 1 > *R_S_*) and other less-invasive means, which sometimes include vaccination, prophylaxis, and social distancing. We find that the incremental benefit of quarantine over symptom monitoring is small for Ebola and SARS, but relatively large for influenza, whose short duration of infectiousness (*D_INF_* ≈ 1-3 days) and some pre-symptomatic infectiousness (*T_0FFSET_* < 0) render symptom monitoring a generally ineffective intervention - particularly in settings with slow contact tracing (*D_CT_* >> 0) and symptom identification (*D_SM_*>> 0). For pandemic influenza strains (which are expected to have higher *R*_0_ than the seasonal influenza strains shown here) or if circumstances arise such that MERS transmissibility increases substantially, quarantine may be necessary to achieve control (**Fig 3B**). Note that CDC quarantine authorities do not extend to pertussis and hepatitis A, and these case studies were selected to demonstrate a broad range of natural histories and transmission routes, including bodily fluid, fecal-oral, and airborne, which will influence the contact networks and traceability of contacts. In general, we find that a reduction in the fraction of contacts who are ultimately traced will decrease the preference for quarantine over symptom monitoring, therefore supporting the previous findings that quarantine was inefficient for a respiratory disease like SARS (20).

Although our results focus on the early stages of an outbreak, contact tracing, symptom monitoring, and quarantine are often key tools for end-stage epidemic control and elimination. As the effective reproductive number decreases below one (e.g. via depletion of susceptible individuals, complementary interventions, seasonality, etc.), our results suggest the preference for quarantine also decreases (**Fig 4**). However, one must consider aspects such as geographic containment, public compliance, and, if the availability of resources lags the epidemic curve, a possible resource-per-case surplus that may enable the more conservative and costly approach of quarantine.

Our results suggest that symptom monitoring could effectively control an outbreak of a new Ebola-like disease, even when infectiousness precedes symptoms and interventions are not perfectly implemented. Because perfect interventions are not always necessary, these results support the conclusion of Cetron et al. (21) that the optimal containment strategy may allow “partial or leaky quarantine” in order to increase the fraction of contacts who participate.

We propose that the most influential parameters should be prioritized for early characterization during an outbreak (22) and should be modeled with conservative consideration of parameter uncertainty, including both real diversity and measurement error. Our framework identifies the key infection-related parameters to define and can form the basis of cost-benefit analyses. Such data-driven decision-making will be critical to determining the optimal public health strategies for the inevitable next epidemic.

## METHODS

### Definitions

“Contact Tracing” is the process of identifying and assessing people who have been exposed to a disease (23). Contacts who are symptomatic when traced are immediately placed in isolation; those who are not symptomatic are placed under either quarantine or symptom monitoring. Here, we model “forward” contact tracing whereby an infected individual names contacts they may have infected (24).

“Isolation” is the separation of a symptomatic individual believed to be infected (23). By reducing the number of risky contact events, isolation reduces disease transmission when infectiousness coincides at least partly with symptoms.

“Quarantine” is the separation of an individual who is believed to be exposed, but is currently not ill (23). If an individual becomes symptomatic, they will be isolated and receive healthcare.

“Symptom Monitoring” is the assessment of symptoms at regular intervals of an individual believed to be exposed, but not ill. If symptoms are detected, the individual is placed in isolation (23). Although they may be encouraged to avoid interpersonal contacts, an individual under symptom monitoring is not separated from others and therefore does not experience a reduction in risky contacts until symptoms are detected.

“Health Seeking Behavior” is the act of seeking healthcare during the presentation of symptoms, leading to isolation. Practically, this intervention could be a health education campaign that prompts individuals to self-identify illness and seek effective isolation. This intervention, which accelerates isolation in a manner separate from contract tracing, provides a comparative care standard for our analysis.

### Model

Individuals in our branching model progress through a Susceptible-Exposed-Infectious-Recovered (SEIR) disease process. We focus our analysis on the early epidemic phase of an emerging infectious disease, assuming no changes to herd immunity within the first few generations of transmission.

Following infection, the number of days before onset of infectiousness and onset of symptoms are the latent period and incubation period, respectively (**Fig 1**). Because clinical symptoms, pathogen concentration, and behavior of the patient can change throughout the course of disease (25), we allow relative infectiousness to vary with time τ since onset of infectiousness (*β*_τ_). The effective reproductive number in the presence of health seeking behavior, symptom monitoring, and quarantine are respectively R_HSB_, R_S_, and R_Q_.

The recent SARS and Ebola epidemics highlighted that hospital isolation does not always contain transmission; we therefore allow isolation effectiveness (γ) to vary to reflect different settings (17, 26, 27). The fraction of contacts who are traced (P_CT_) can be less than 1, encompassing symptomatic infectors who fail to recall contacts, asymptomatic “silent” infection events, reluctance to report contacts, and challenges in identifying contacts, especially for airborne transmission routes. Imperfections and uncertainties in risk profiling can reduce the fraction of traced contacts that are truly infected (P_INF_) (16, 20). Delays in tracing a contact (D_CT_) can arise for numerous reasons, including intractable roads, low mobile phone penetration, and personnel limitations. The delay between symptom onset and isolation (D_SM_) specifically applies to individuals under symptom monitoring and is influenced by the frequency of monitoring, delays in recognizing sometimes unreliable clinical features, and delays in prompt isolation upon symptom detection. Intervention performance parameter examples can be found in **Table 1**.

### Simulation

We draw disease characteristics for each simulated individual from disease-specific input distributions. During each hour τ of infectiousness, an individual infects a number of new individuals drawn from a Poisson distribution (or, if super-spreading factor *κ* ≠ 1, a negative binomial distribution (18)) with mean equal to the product of the expected number of onward infections for the individual (*R*_0_) and the relative infectiousness *β*_τ_ where 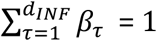. We assume time-varying relative infectiousness follows a triangular distribution with time of peak infectiousness (τ_*β*_) occurring anywhere between the onset and end of infectiousness, inclusively.

We record both the day of transmission and the source of each new infection for each transmission event, and draw disease characteristics for each newly infected individual. An individual is identified by contact tracing with probability P_CT_ at the earlier time of either when their infector is isolated or the time of the transmission event if infection occurs while the infector was isolated. After an operational lag time of D_CT_ days, a contact is placed under quarantine, symptom monitoring or, if already symptomatic, isolation. An individual in isolation or quarantine has their infectiousness reduced by a factor γ for the remainder of their disease. An individual under symptom monitoring is isolated D_SM_ days after symptom onset. A full description of the model process can be found in **S1 Appendix**.

### Parameterization

As compared to characteristics related to the natural history of symptoms and illness, key aspects of the natural history of infectiousness tend to be harder to observe and measure (28). Therefore, we use a Sequential Monte Carlo particle filtering algorithm (29, 30) to create a joint probability space of the time offset between the latent period and incubation period (*T_0PPSET_* = *T_LAT_* – *T_INC_*), time of peak infectiousness (τ_*β*_), and duration of infectiousness (*d_INP_*). From an uninformative prior distribution of each parameter bounded by published observations, we simulate five infection generations of 500 initial individuals and record the simulated serial interval (i.e., the time between symptom onset in infector-infectee pairs). Parameter sets are resampled with importance weights determined by the degree to which the distribution of simulated serial intervals match published serial interval distributions, using the Kolmogorov-Smirnov test of the difference between cumulative distribution functions (**Table 2**) (31, 32). After perturbation, the process is repeated until convergence, which we define to be when the median Kolmogorov-Smirnov statistic was within 10% of the previous two iterations. This cutoff criterion balances the objectives of finding a stationary posterior set of particles while preserving some of the heterogeneity in input parameters.

Holding the incubation period distribution constant, we fit an offset for the latent period (*T_0FFSET_*) for several reasons, including consistency with CDC methods for disease characterization (33), the biological expectation of these timings both being linked to pathogen load, and to parsimoniously limit each characteristic to one interpretable parameter. For the duration of infectiousness (*d_INF_*), we fit the upper bound of a uniform distribution with a lower bound of 1 day. To allow for variable infectiousness during this duration, we assume a triangular distribution of relative infectiousness *β*_τ_ and fit the time of peak infectiousness (τ_*β*_). A full description of the model parameterization can be found in the **S2 Appendix.**

### Analysis

Partial rank correlation coefficients are calculated to identify the most influential disease characteristics (e.g. duration of infectiousness) and intervention performance metrics (e.g. isolation effectiveness). In order to remove dependence between the parameters jointly fit through the particle filtering method above, we use Latin Hypercube Sampling to draw 5,000 sets from each marginal posterior parameter distribution independently. To maximize coverage of the parameter space we allowed fractional parameters (γ, P_CT_, P_INF_, *k*) to range from 0 to 1, delays (D_CT_, D_SM_) to range from 0 to 7 days, *R_0_* to range from 1 to 5, and the incubation period (*T_INC_*) to be shrunk by up to 50% or stretched by up to 150%.

We draw 1,000 samples from the joint-parameter space from the particle filtering method and measure R_0_, R_Q_, R_S_, and R_HSB_ for each disease. We compare the effectiveness of symptom monitoring and quarantine by the absolute difference (*R_S_* – *R_Q_*) and the relative difference 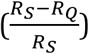. We calculate the number of days an infected individual was in quarantine but not yet infectious (*d_Q_*) as surrogate for the marginal cost of quarantine over symptom monitoring. As abstract surrogates for cost-effectiveness, we calculate the absolute difference per quarantine day (*R_S_* – *R_Q_*)/*d_Q_* and relative difference per quarantine day 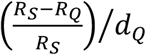 and present these results in Figs S1 and S3. More concrete measures of cost-effectiveness would require economic and social considerations that are outside the scope of this paper.

When risk profiling is imperfect (i.e. P_INF_ < 1), uninfected individuals may be mistakenly traced as contacts and placed under symptom monitoring or quarantine. Such events may be conceptualized as false positives and will decrease P_INF_; conversely, individuals who are infected but not traced are false negatives and will decrease P_CT_. We assume that non-infected contacts are followed for a length of time set up the 95^th^ percentile incubation period 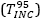, at which point health authorities may conclude the contact was not infected after all. This changes the number of days in quarantine to

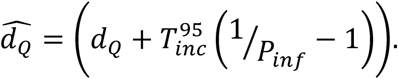

## ACKNOWLEDGEMENTS

CMP, LMC, and COB were supported by Cooperative Agreement U54GM088558 from the National Institute Of General Medical Sciences. CMP was also supported by National Research Service Award T32AI007535-16A1. YHG was supported by the National Institutes of Health (K08-AI104767), the Smith Family Foundation, and the Doris Duke Foundation. The content is solely the responsibility of the authors and does not necessarily represent the official views of the National Institute Of General Medical Sciences or the National Institutes of Health. The funders had no role in study design, data collection and analysis, decision to publish, or preparation of the manuscript. The authors thank Colin Worby for helpful statistical discussions.

## S1 Appendix. Disease model

The model simulates a branching network of infected individuals only. An individual *i* is assigned characteristics sampled from distributions defined for each disease (**Table S1**). The incubation period (*T_INC_*), i.e. the time from infection to symptom onset, is drawn from published distributions (**Table 2**). The duration of infectiousness (*d_INF_*), time of peak infectiousness (τ_*β*_), and time offset between the ends of the latent and incubation periods (*T_OFFSET_*) are drawn from the joint posterior distribution generated by the sequential Monte-Carlo (SMC) particle filtering method described in **S2 Appendix**. For clarity, we describe the method for an individual *i*, but the following process is repeated for an initial population of 1,000 individuals who each initiate distinct trees.

The expected number of onward infections by individual *i, R*_0*i*_., is distributed over each hour τ of disease *R_τi_* = *β_τi_.R*_0*i*_., where *β_τi_* is the relative infectiousness of individual *i* on hour τ such that 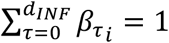 (**Fig S6**). For parsimony and ease of interpretation, we assume *β_τi_* follows a discretized triangle distribution with a peak value at time *τ_β_* drawn from the SMC posterior and rounded to the nearest hour. When 0 < *τ_β_* < *d_INF_, β_τi_*. is defined by a piecewise function according to whether τ is in the period before the peak of infection (*τ* ≤ *τ_β_*) or τ is after the peak of infection:

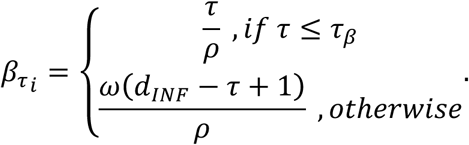

Infectiousness after the peak is scaled by *ω*, which is the slope of the post-peak infectiousness line:

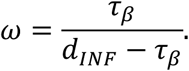

In order to normalize *β_τi_* to sum to 1, each piecewise component is divided by *ρ*, which is defined as the sum of infectiousness before the peak and after the peak:

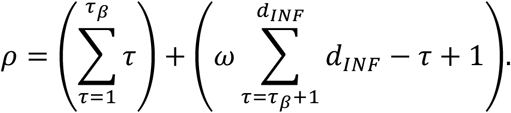

Under the simple conditions of linearly increasing or decreasing infectiousness, *β_τi_* is respectively defined by

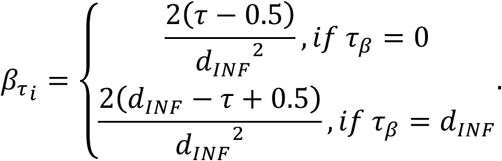

The total number of infections (*N_i_*) generated by individual *i* is drawn from a negative binomial distribution with mean equal to the total expected number of infections (*E*(*N_i_* = *R*_0*i*_) generated by individual *i* and dispersion factor *k*. If *k* = 1, the negative binomial distribution reduces to a Poisson distribution with rate *λ* = *R*_0*i*_. If *k* < 1, the number of infections generated per case will be overdispersed to simulate super-spreading (**Fig S5**).

If individual *i* generates infections (i.e., *N_i_* > 0), then each infection generated is assigned to a particular time τ drawn from a random sample *τ ∊* {0,1,…, *d_INF_*_*i*_} that is weighted by *R*_τ__i_/*R*_0*i*_ so that hours of high infectiousness are more likely to have larger values of *N*_τi_ Therefore, each individual *i* has a vector of 〈*N*_0*i*_, *N*_1*i*_,…, *N_d_inf_i___*〉 of onward infections that occur during each hour *τ* of their infectiousness.

A new individual *j* is generated for each onward infection *N_t i_* ≥ 1. Disease characteristics for individual *j* are drawn as above and are set to begin at the time of infection of individual *j*.

Each individual *j* will be traced with probability *P_CT_.* If traced, individual *j* is placed under symptom monitoring or quarantine after an operational lag time of *D_CT_* days. The lag time occurs after the earlier of: (1) isolation of individual *i* or (2) removal of individual *i* from the disease system upon recovery or death. We assume individuals granted access to the quarantine or isolation room will be logged and given the same attention as contacts traced through epidemiological interview and will therefore will begin monitoring or quarantine at the time of the infection control breach or transmission event.

Next we determine the time of isolation for individual *j*. If time of symptom onset for individual *j* occurs before individual *j* is traced, individual *j* is immediately isolated. Otherwise, time of isolation for individual *j* depends on whether symptom monitoring or quarantine is used. Under symptom monitoring, isolation of individual *j* occurs a delay *D_SM_* days after symptom onset. Note that for contacts checked twice-daily, *D_SM_*~unif(0,0.5). Upon isolation, the hourly expected number of onward infections is reduced to *R_τ _j__* = (1 – *γ*)*β_τ_j__ R*_0_*j*__ where *γ* is effectiveness of isolation with support [0,1]. If individual *j* is under quarantine, then *R_t_j__* is reduced by (1 – *γ*) beginning at the time individual *j* is traced.

## S2 Appendix. Parameterization via Sequential Monte Carlo

We used the following method to generate parameter sets that are consistent with the published serial interval and incubation period distributions for each case study disease. The objective of this procedure is not explicitly parameter inference, for which raw data and disease-specific nuance is necessary, but rather a range of parameters to reflect the heterogeneity of each disease in a common framework. Data-informed incubation period and serial interval distributions were collected through a literature review. Sequential Monte-Carlo (also known as particle filtering) methods were used to estimate the joint distribution of three disease parameters (*T_0FFSET_, d_INF_,* and *τ_β_*) using knowledge of the incubation period and serial interval distributions [28,29].

Here we assume that the latent period for an individual ends some time (*T_OFFSET_*) before (T_OFFSET_ < 0) or after (T_OFFSET_ > 0) the onset of symptoms. Therefore, *T_0FFSET_* is a translation of the incubation period distribution. We assume a uniform distribution of duration of infectiousness from 1 day to *d_INF_*, by hour. We assume the distribution of relative infectiousness to follow a triangle distribution with a peak at time *τ_β_*, which ranges from 0 (indicating infectiousness is linearly decreasing) to 1 (indicating infectiousness is linearly increasing).

The steps are as follows:

i. Use the “lhs” package in R to create a Latin Hypercube sample of 1000 parameter sets *Θ*_1_, *Θ_2_,*…, *Θ*_1000_ consisting of *T_OFFSET_, d_INF_*, and *τ_β_*, which are bounded by the range of the parameter values found in the literature (**Table 2**)
ii. Draw a parameter set *Θ_i_* = *Θ*_1_
iii. Under parameter set *Θ_i_,* run the branching epidemic model beginning with 500 distinct infections under a situation with no interventions.
iv. Use the two-sample “ks.test” function in R to calculate the Kolmogorov-Smirno (*KS_i_*) test statistic comparing the empirical distribution of simulated serial intervals to the published serial interval distribution (**Fig S4**).
v. Repeat steps (iii-iv) for each parameter set *Θ*_1_,= *Θ_2_, Θ_3_,*…, *Θ*_1000_.
vi. Set an adaptive threshold KS* equal to 80% of the maximum *KS_t_* among the 1000 parameter sets above.
vii. Create a set of 1000 Θ_candidate_ parameter sets by selecting a weighted sample with replacement from all parameter sets *Θ_i_* with KS_i_ ≤ KS*. Weights are proportional to 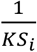.
viii. Perturb each Θ_candidate_ by between 0% and 2% (uniformly) of the initial parameter value range.
ix. Repeat steps (ii)-(viii) until the median KS is within 10% of each of the previous two rounds.

It is widely known that generation intervals are difficult to observe in nature, but challenges also arise in measurements of generation intervals in simulations. For example, generation time distributions may change over the course of an epidemic due to depletion of the susceptible individuals (1). Analogous to the approach of (2), our method of direct observation of simulated serial intervals is restricted to the exponential growth period of an epidemic, as produced by a branching model. However, the potential for the length of generation intervals to be under-estimated near the peak of an epidemic can cause a downward bias in the published serial intervals upon which our parameterization methods are based (2, 3). Therefore, this downward bias can result in a bias towards “faster” disease parameters (namely, a leftward bias in *T_0FFSET_, d_INF_,* and *τ_β_*). Such a bias would reduce the simulated effectiveness of all interventions considered in this paper.

---

**Fig. S1.**
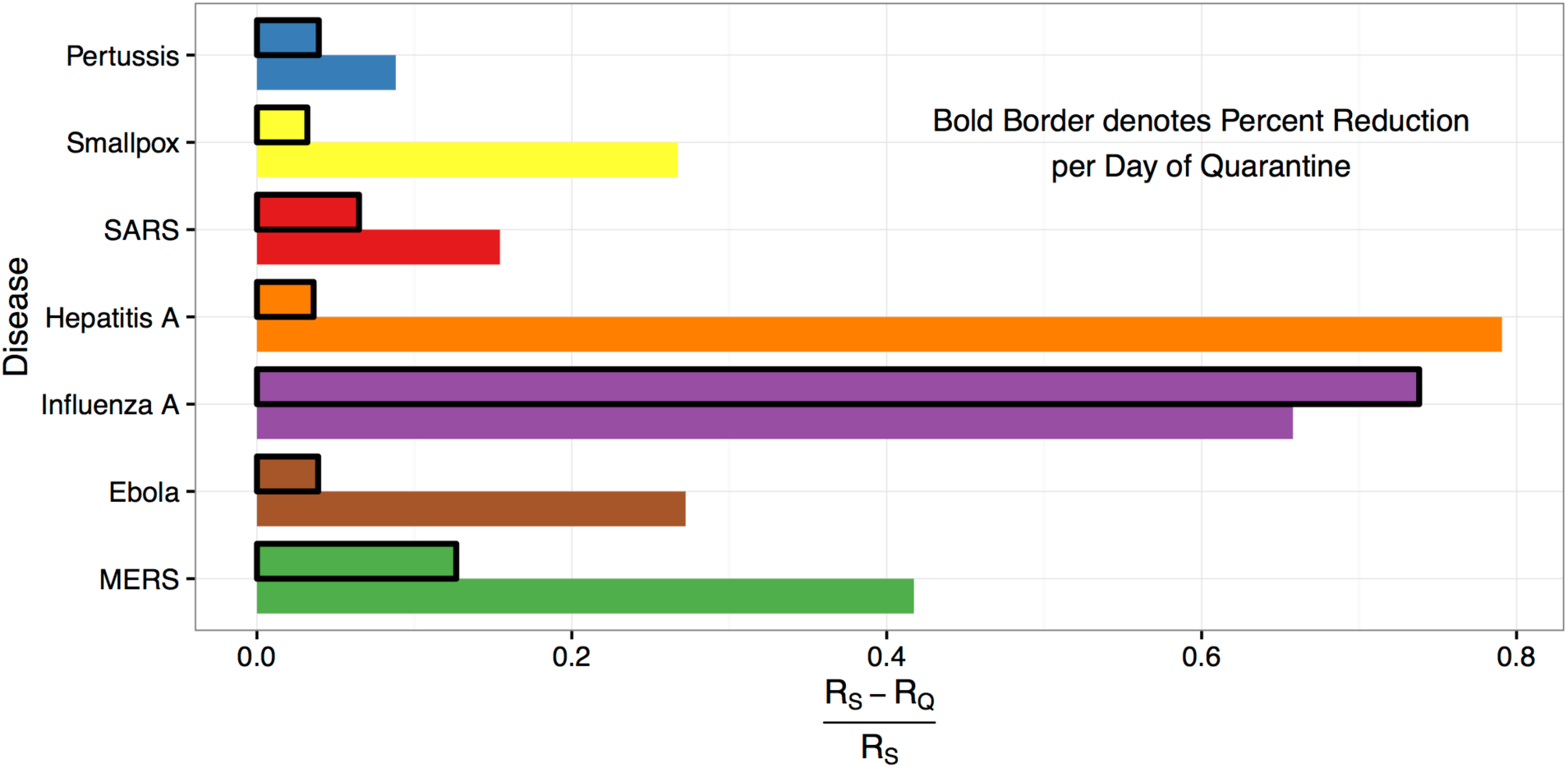
Relative comparative effectiveness and cost-effectiveness. The relative comparative effectiveness varies widely by disease, with quarantine reducing *R_S_* by >65% for influenza A and hepatitis A and by <10% for pertussis. However, due to a much shorter incubation period of influenza A versus hepatitis A (**Table 2**), the relative cost-effectiveness measured by the reduction per day of quarantine (outlined bars) is substantially higher for influenza A than hepatitis A.

**Fig. S2.**
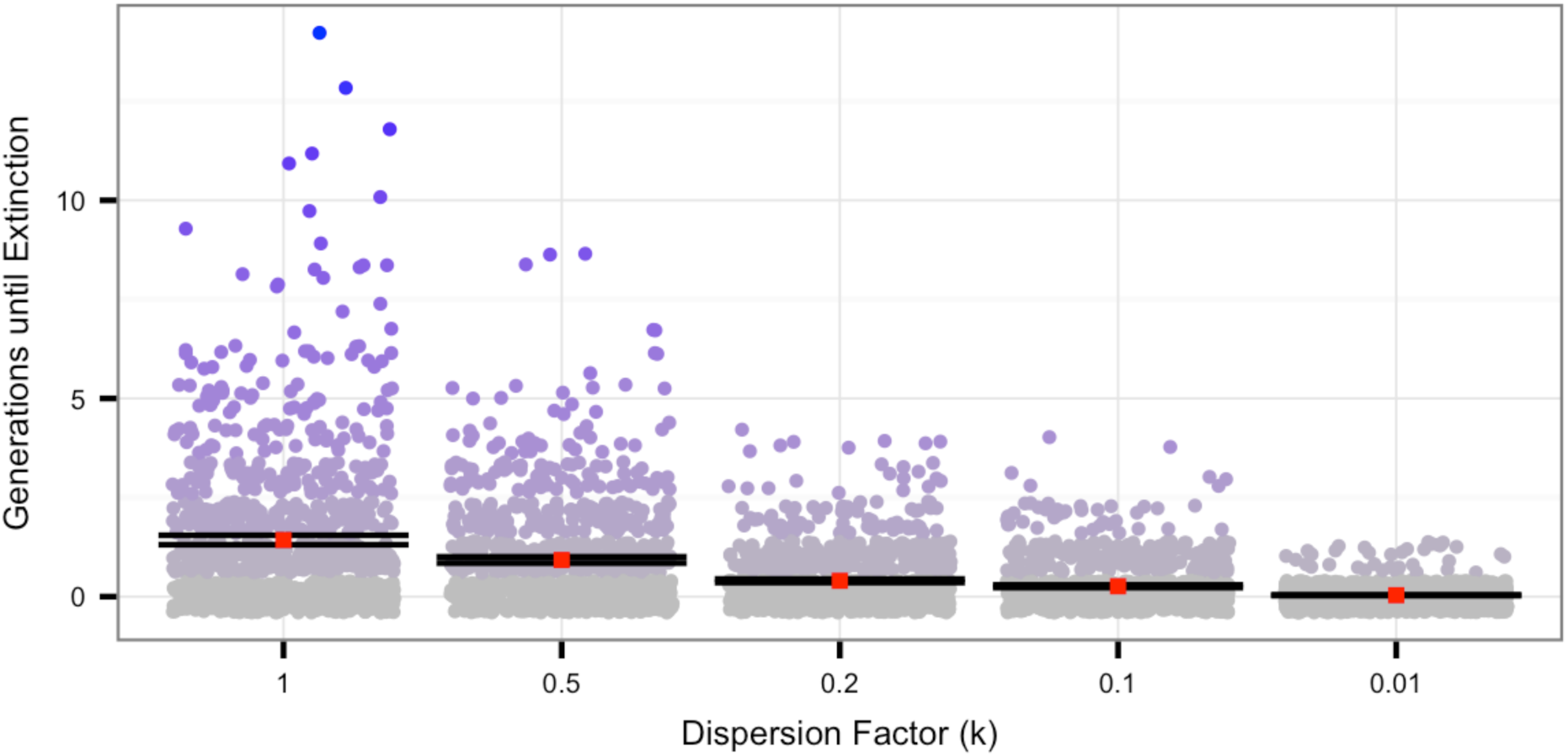
Number of generations until extinction decreases in the presence of overdispersion. As the dispersion factor (k) decreases (i.e., creating more super-spreading), the average number of generations (red square) before extinction of a single infectious case with an R_0_ of 0.75 is 1.43 for k=1, 0.26 for k=0.1, and 0.036 for k=0.01. Each point indicates the number of generations (y axis, jittered for visibility) until a transmission tree initiated by a single case is contained. 95% confidence intervals shown in brackets.

**Fig. S3.**
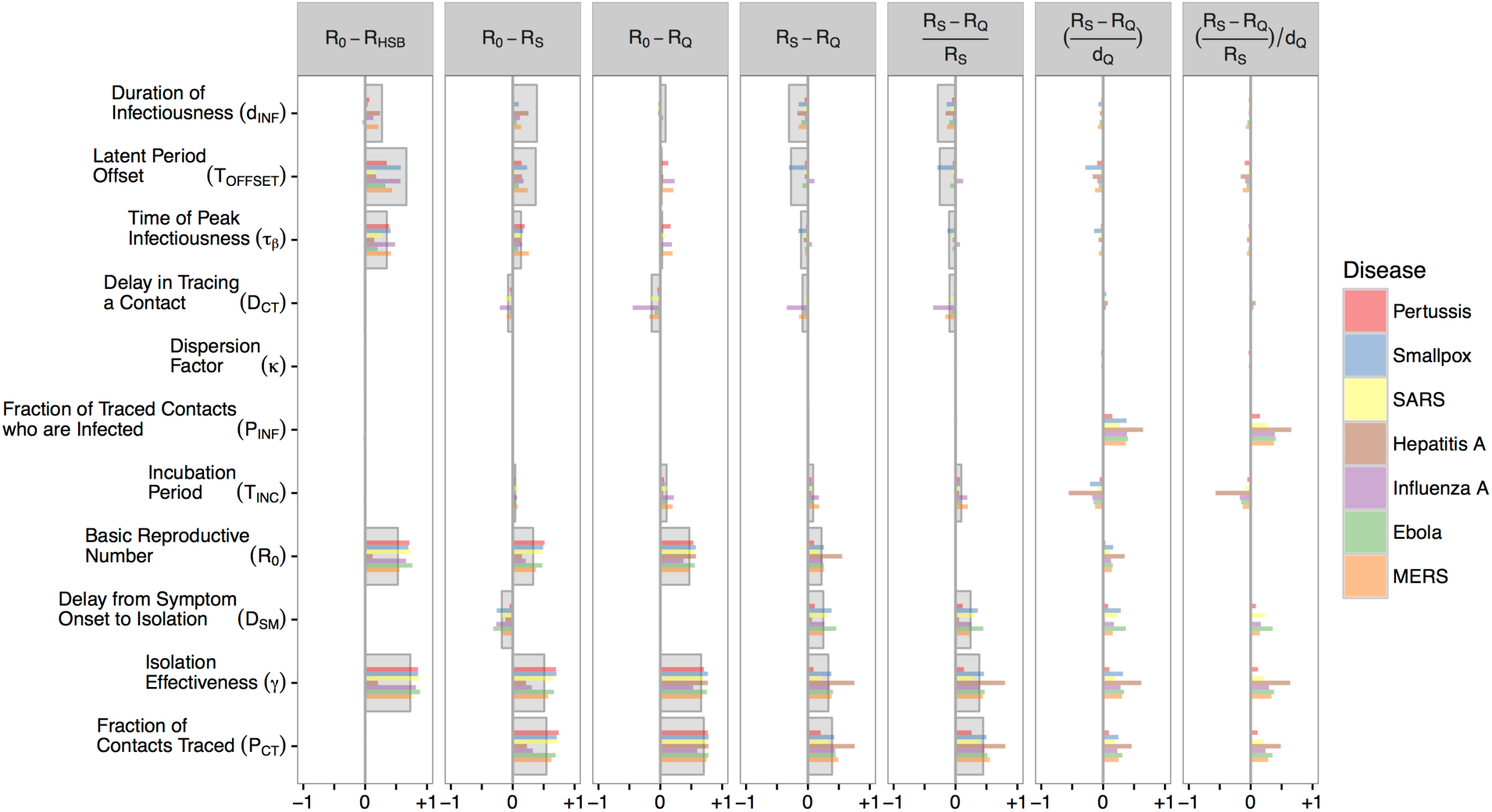
Partial rank correlation coefficients for all outcomes. Partial rank correlation coefficients (x-axis) measuring the influence of disease characteristics and intervention performance metrics (rows) on the impact, comparative effectiveness, and comparative cost-effectiveness of the interventions under study. Disease-specific estimates are shown with colored bars and pooled estimates with larger grey bars. For example, increasing the delay in tracing a named contact D_CT_ has a generally small effect negative effect on R_S_-R_Q_ when pooled across diseases (large grey bar), but for influenza A specifically (purple bar), D_CT_ has a rather large negative effect on R_S_-R_Q_. Note that pooled estimates for comparative cost-effectiveness are not available due to non-monotonic relationships across diseases.

**Fig. S4.**
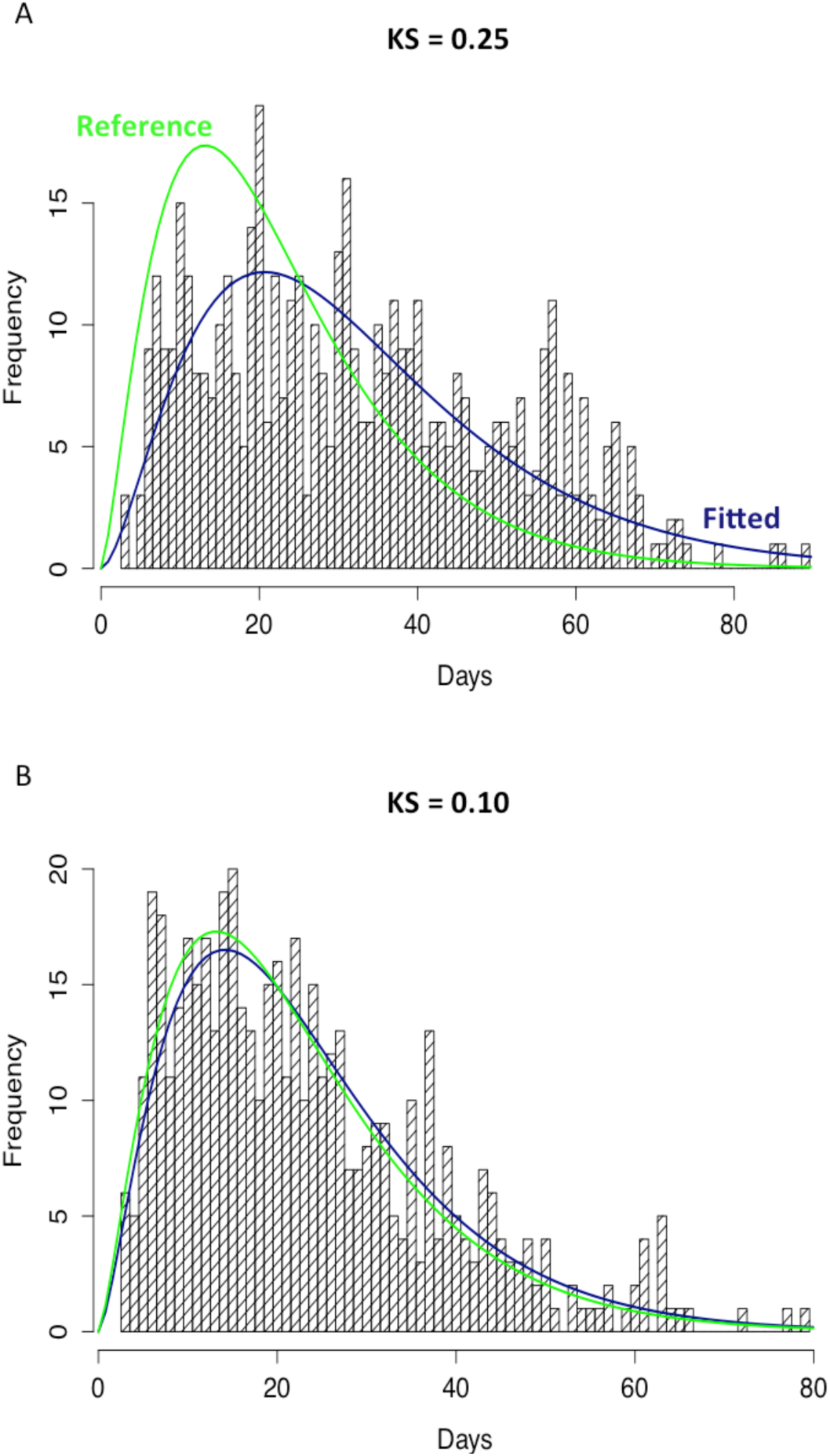
Demonstration of the Kolmogorov-Smirnov distance for SMC paramterization method. The parameter set in Panel A generates serial intervals (bars, blue line) that are poorly explained by the reference serial interval distribution (green line), generating a KS score of 0.25. A later iteration of the SMC algorithm generated parameter set (B) in which the generated serial intervals are more consistent with the reference serial interval (same green line).

**Fig. S5.**
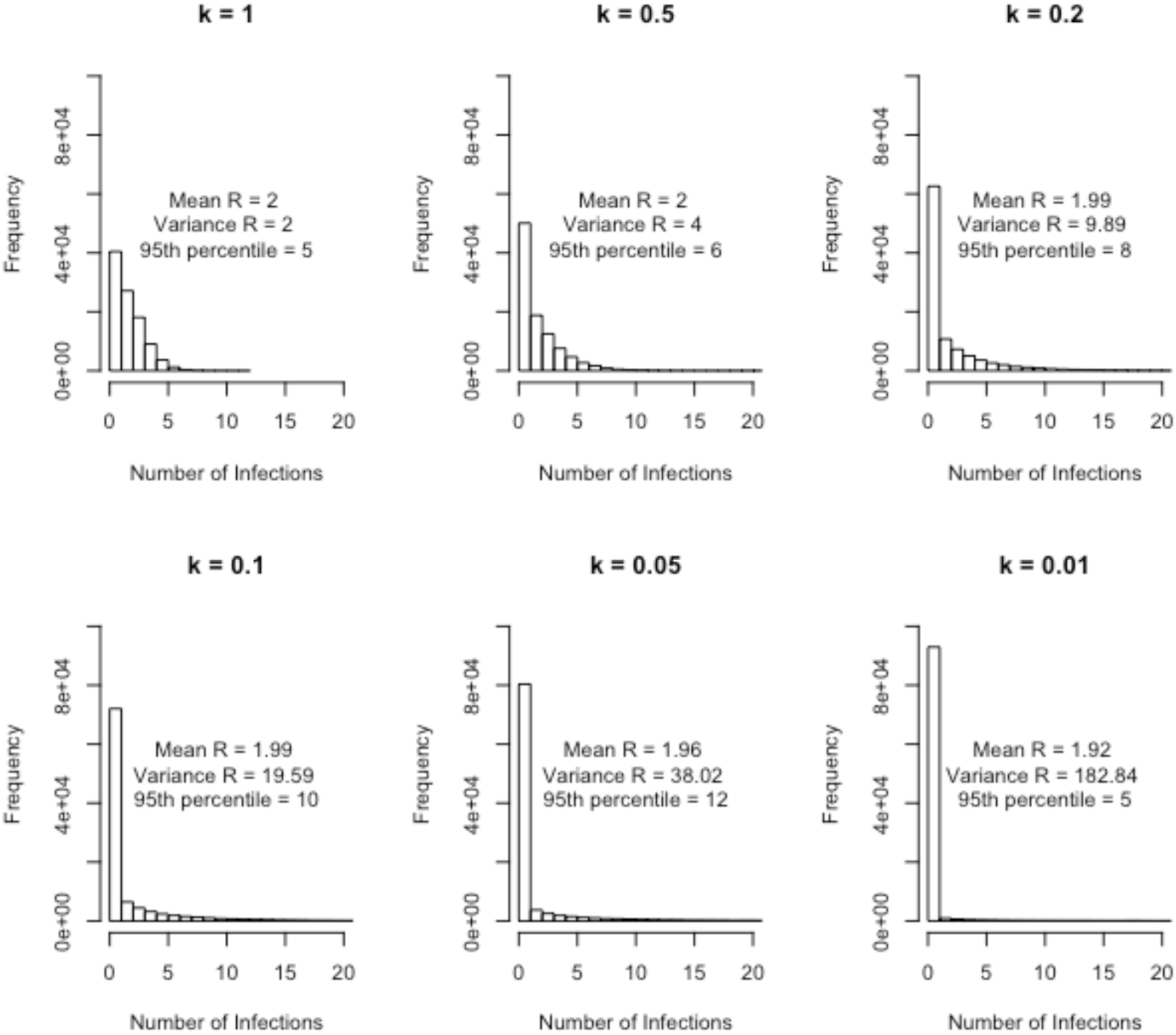
Demonstration of overdispersion parameter (k). In a system with 100,000 agents, as the dispersion parameter (k) decreases (i.e., creating more super-spreading), the variance of the number of infections generated by each infectious individual increases while the mean is approximately constant at the input value of 2.

**Fig. S6.**
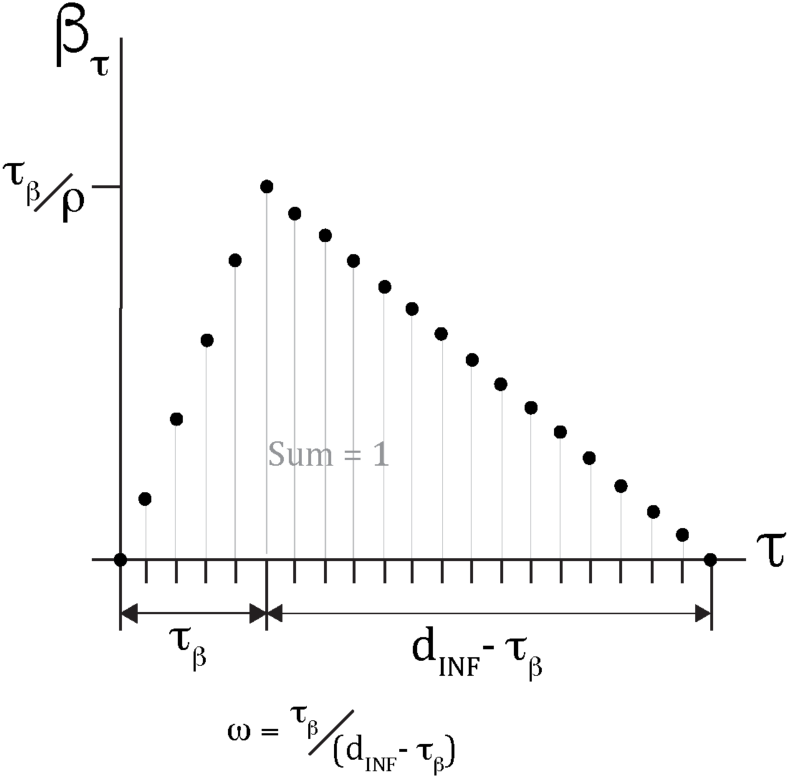
Distribution of Infectiousness. Key time points in the distribution of infectiousness include the peak (*τ_β_*) and duration (*d_INF_*) of infectiousness.

## REFERENCES

1. McMichael AJ (2004) Environmental and social influences on emerging infectious diseases: past, present and future. Philos Trans RSocB Biol Sci 359(1447):1049–1058.

2. Morens DM, Folkers GK, Fauci AS (2004) The challenge of emerging and re-emerging infectious diseases. Nature 430.

3. Jones KE, Patel NG, Levy MA (2008) Global trends in emerging infectious diseases. Nature 451(7181):990–993.

4. Barbera J, et al. (2001) Large-Scale Quarantine Following Biological Terrorism in the United States. J Am Med Assoc 286(21):2711–2717.

5. American Civil Liberties Union, Yale Global Health Justice Partnership (2015) Fear, Politics, and Ebola. How Quarantines Hurt the Fight Against Ebola and Violate the Constitution Available at: https://www.aclu.org/sites/default/files/field_document/aclu-ebolareport.pdf.

6. Médecins Sans Frontières (MSF) (2014) Ebola: Quarantine can undermine efforts to curb epidemic. Available at: http://www.msf.org/article/ebola-quarantine-can-undermine-efforts-curb-epidemic.

7. Drazen JM, et al. (2014) Ebola and Quarantine. NEngl JMed 371(21):2029–30.

8. Day T, Park A, Madras N, Gumel A, Wu J (2006) When Is Quarantine a Useful Control Strategy for Emerging Infectious Diseases? Am J Epidemiol 163(5):479–485.

9. Gates B (2015) The Next Epidemic – Lessons from Ebola. 1–4.

10. Heymann DL ed. (2014) Control of Communicable Diseases Manual (APHA Press). 20th Editi.

11. World Health Organization (2005) International Health Regulations Available at: http://apps.who.int/iris/bitstream/10665/43883/1/9789241580410_eng.pdf.

12. Centers for Disease Control and Prevention (2014) Legal Authorities for Isolation and Quarantine Available at: http://www.cdc.gov/quarantine/pdf/legal-authorities-isolation-quarantine.pdf.

13. Rothstein MA, et al. (2003) Quarantine and isolation: Lessons learned from SARS Available at: http://www.iaclea.org/members/pdfs/SARSREPORT.Rothstein.pdf.

14. Nyenswah T, et al. (2015) Controlling the Last Known Cluster of Ebola Virus Disease – Liberia, January - February 2015. 64(18):500–504.

15. CDC (2014) Interim U.S. Guidance for Monitoring and Movement of Persons with Potential Ebola Virus Exposure (Atlanta, Georgia) Available at: http://www.cdc.gov/vhf/ebola/exposure/monitoring-and-movement-of-persons-with-exposure.html.

16. Chung WM, et al. (2015) Active Tracing and Monitoring of Contacts Associated With the First Cluster of Ebola in the United States. Ann Intern Med (163):164–173.

17. Fraser C, Riley> S, Anderson> RM, Ferguson NM (2004) Factors that make an infectious disease outbreak controllable. Proc Natl Acad Sci 101:6146–6151.

18. Lloyd-Smith JO, Schreiber SJ, Kopp PE, Getz WM (2005) Superspreading and the effect of individual variation on disease emergence. Nature 438(7066):355–359.

19. Gostic KM, Kucharski AJ, Lloyd-Smith JO (2015) Effectiveness of traveller screening for emerging pathogens is shaped by epidemiology and natural history of infection. Elife 2015 (4):1–16.

20. Glasser JW, Hupert N, McCauley MM, Hatchett R (2011) Modeling and public health emergency responses: Lessons from SARS. Epidemics 3(1):32–37.

21. Cetron M, Maloney S, Koppaka R, Simone P (2004) Isolation and Quarantine: Containment Strategies for SARS 2003. Institute of Medicine (US) Forum on Microbial Threats, eds Knobler S, Mahmoud A, Lemon S, pp 71–83.

22. Lessler J, Edmunds WJ, Halloran ME, Hollingsworth TD, Lloyd AL (2015) Seven challenges for model-driven data collection in experimental and observational studies. Epidemics 10:78–82.

23. Centers for Disease Control and Prevention (2014) CDC Methods for Implementing and Managing Contact Tracing for Ebola Virus Disease in Less-Affected Countries.

24. Muller J, Kretzschmar M, Dietz K (2000) Contact tracing in stochastic and deterministic epidemic models. Math Biosci 164:39–64.

25. Fine P, Eames K, Heymann DL (2011) “Herd immunity”: A rough guide. Clin Infect Dis 52(7):911–916.

26. Klinkenberg D, Fraser C, Heesterbeek H (2006) The effectiveness of contact tracing in emerging epidemics. PLoS One 1(1):e12.

27. Lloyd-Smith JO, Galvani AP, Getz WM (2003) Curtailing transmission of severe acute respiratory syndrome within a community and its hospital. Proc Biol Sci 270(1528):1979–1989.

28. Richardson M, Elliman D, Maguire H, Simpson> J, Nicoll A (2001) Evidence base of incubation periods, periods of infectiousness and exclusion policies for the control of communicable diseases in schools and preschools. Pediatr Infect Dis J 20(4):380–91.

29. Andrieu C, Doucet A, Singh SS, Tadic VB (2004) Particle methods for change detection, system identification, and control. Proc IEEE 92(3):423–438.

30. Doucet A, de Freitas N, Gordon N (2001) An Introduction to Sequential Monte Carlo Methods. Seq Monte Carlo Methods Pract:3–14.

31. Djuric PM, Miguez J (2010) Assessment of Nonlinear Dynamic Models by Kolmogorov-Smirnov Statistics. IEEE Trans Signal Process 58(10):5069-5079.

32. Marsaglia G, Tsang WW, Wang J (2003) Evaluating Kolmogorovs Distribution. J Stat Softw 8(18):1–4.

33. Centers for Disease Control and Prevention (2015) Epidemiology and Prevention of Vaccine Preventable Diseases eds Hamborsky J, Kroger A, Wolfe C (Public Health Foundation, Washington D.C.). 13th ed.

## Supplemental References

1. Scalia Tomba G, Svensson Å, Asikainen T, Giesecke J (2010) Some model based considerations on observing generation times for communicable diseases. Math Biosci 223(1):24–31.

2. Svensson Å (2007) A note on generation times in epidemic models. Math Biosci 208(1):300–311.

3. Kenah E, Lipsitch M, Robins JM (2008) Generation interval contraction and epidemic data analysis. Math Biosci 213(1):71–9.

